# A community-based approach to image analysis of cells, tissues and tumors

**DOI:** 10.1101/2021.07.22.451363

**Authors:** CSBC/PS-ON Image Analysis Working Group, Juan Carlos Vizcarra, Erik A. Burlingame, Clemens B. Hug, Yury Goltsev, Brian S. White, Darren R. Tyson, Artem Sokolov

**Author notes:** Consortia authors listed in Supplemental Table 1. The Jackson Laboratory for Genomic Medicine, Farmington, CT, USA. These authors contributed equally (Brian S. White, Darren R. Tyson, Artem Sokolov).

## Abstract

Emerging multiplexed imaging platforms provide an unprecedented view of an increasing number of molecular markers at subcellular resolution and the dynamic evolution of tumor cellular composition. As such, they are capable of elucidating cell-to-cell interactions within the tumor microenvironment that impact clinical outcome and therapeutic response. However, the rapid development of these platforms has far outpaced the computational methods for processing and analyzing the data they generate. While being technologically disparate, all imaging assays share many computational requirements for post-collection data processing. We convened a workshop to characterize these shared computational challenges and a follow-up hackathon to implement solutions for a selected subset of them. Here, we delineate these areas that reflect major axes of research within the field, including image registration, segmentation of cells and subcellular structures, and identification of cell types from their morphology. We further describe the logistical organization of these events, believing our lessons learned can aid others in uniting the imaging community around self-identified topics of mutual interest, in designing and implementing operational procedures to address those topics and in mitigating issues inherent in image analysis (e.g., sharing exemplar images of large datasets and disseminating baseline solutions to hackathon challenges through open-source code repositories).

## 1 Introduction

The spatial organization and dynamic interactions of cells in a tumor microenvironment profoundly impact cancer clinical outcomes and therapeutic responses. Researchers within the Cancer Systems Biology Consortium (CSBC) and the Physical Sciences - Oncology Network (PS-ON) are actively interrogating these interactions by highly-multiplexed and/or time-resolved, subcellular resolution imaging platforms. However, the rapid development of the imaging platforms has far outpaced the computational methods for processing and analyzing the data they generate, in part because method development is often borne independently and repeated by individual research groups. To accelerate the development of computational methods and close the gap between data collection and analysis, the CSBC/PS-ON Image Analysis Working Group (IAWG) has been focused on consolidation of development efforts across research groups and effective dissemination of image analysis tools and ideas across the CSBC/PS-ON centers and with external consortia, including the Human Tumor Atlas Network (HTAN) and Human BioMolecular Atlas Program (HuBMAP). In high-dimensional digital pathology, dozens of spatially resolved molecular markers are collected from millions of cells per specimen, providing an unprecedented view of single cells in the setting of an intact tissue. Technologies such as cyclic immunofluorescence (CyCIF) (Lin et al., 2018), co-detection by indexing (CODEX) (Goltsev et al., 2018), imaging mass cytometry (IMC) (Giesen et al., 2014), and multiplexed ion beam imaging (MIBI) (Angelo et al., 2014) measure co-localized abundance of 50–100 proteins and protein modifications. Likewise, live cell tracking experiments characterizing scores of unique conditions can include hundreds of cells imaged in multiple channels every 6–20 minutes over several days (Neumann et al., 2010; Quaranta et al., 2009; Tyson et al., 2012). These modern approaches yield orders of magnitude more data than traditional haematoxylin and eosin (H&E) and immunohistochemistry (IHC) staining. For example, a single canonical whole-slide image produced by CyCIF is on the order of tens of gigabytes, which raises new challenges for data storage, processing and analysis that were not critical for the more traditional imaging methods.

To unlock potential biological or clinical insight, the analysis software for highly multiplexed and/or time-resolved images must address a range of image processing tasks. A canonical workflow includes image stitching and registration, illumination correction, cell and nuclear segmentation and/or tracking, assignment of cell type identity, and recognition of high-level spatial or temporal features that may be characteristic of disease phenotypes. Each step comes with its own set of challenges, and lessons learned by one research group are not always effectively communicated to other groups, even within the same consortium. Software engineering challenges also abound, with open-source software designed around specific image processing steps (e.g., segmentation) but with little consideration for interoperability or integration within a larger end-to-end pipeline framework. This is in stark contrast to sequence data—both single-cell and bulk modalities—for which well-established frameworks, such as GATK (Auwera and O’Connor, 2020), Seurat (Satija et al., 2015) and Galaxy (Afgan et al., 2018), effectively combine software modules for processing a raw sequence file through alignment, quantification, differential gene expression and pathway enrichment to identify potential associations with a phenotype of interest.

Our objective was to identify and begin to address impediments to image analysis shared by the biomedical research community. To achieve this, we hosted a two-day workshop in January, 2020 in Seattle, WA, where we invited members of CSBC/PS-ON centers and external speakers to highlight challenges that hold particular relevance to their work. A subset of the presented challenges was selected to be addressed in a hackathon held in March, 2020 in Nashville, TN.

To our knowledge, this is the first effort to address multiple aspects of the canonical image analysis workflow within a hackathon format. While all hackathons involve some amount of preliminary planning and organization (Ahmed et al., 2018; Connor et al., 2019; Fecho et al., 2019; Ferreira et al., 2019; hackseq Organizing Committee 2016, 2017), the hosting of a formal face-to-face workshop has allowed us to not only define hackathon challenges, but also formalize the larger workflow connecting them. Unlike previous image analysis hackathons that focused on further development and application of specific tools, e.g., 3D Slicer (Kapur et al., 2016) and Fiji (Schindelin et al., 2012), we allowed the participants to utilize any existing methods and encouraged the development of new ones. Our approach also differed from online competitions such as the Kaggle 2018 Data Science Bowl on nuclear segmentation (Caicedo et al., 2019) and the Cell Tracking Challenge (Ulman et al., 2017), in which teams work remotely and benefit from relatively long time periods to solve one specific task. Instead, we brought together researchers at one physical location, which furthered our additional consortia-wide goals of providing an educational experience to trainees, spurring collaboration within the consortia, and disseminating research perspectives across diverse backgrounds. Here, we summarize our logistical efforts and the resulting output of the 2020 workshop and hackathon, as well as provide our perspective for the future of collaborative large-scale image analysis.

## 2 Methods

### 2.1 Image analysis working group

The IAWG is a joint forum within which CSBC and PS-ON scientists, as well as external invited speakers, share their image analysis and visualization results and discuss open research questions. These questions reflect the diversity of biological domains and imaging modalities of CSBC and PS-ON. Researchers in CSBC employ imaging techniques (e.g., CyCIF, CODEX, MIBI, multiplexed IHC, and time-lapse fluorescence microscopy) to study biological phenomena such as tumor-immune interactions and the tumor microenvironment, drug resistance/sensitivity, metastasis, and tumor heterogeneity across many cancer types. Often, the imaging data are integrated with other widely-used systems biology methods, including sequencing, mechanistic modeling, machine learning, evolution / ecology, and network inference. In contrast, researchers in PS-ON combine imaging with approaches from the physical and mathematical sciences to study mechanical cues, transport phenomena, bioelectric signals, thermal fluctuations, and spatio-temporal organization of cancer at scales ranging from subcellular to organ and whole organism. CSBC and PS-ON share a coordinating center that facilitates collaboration (including support for the IAWG workshop and hackathon), resource sharing, and education and outreach across the two consortia and with the external scientific community.

### 2.2 Workshop

#### 2.2.1 Pre-workshop planning and funding

Initial monthly presentations of the IAWG revealed a substantial overlap in image analysis interests and challenges shared by CSBC and PS-ON researchers. This motivated us to host a workshop oriented around computational image analysis challenges that could realistically be addressed in a subsequent two-day hackathon. Because we anticipated that such challenges would transcend biological domains and specific imaging modalities, we advertised the workshop within the CSBC/PS-ON community and broadly in other consortia leveraging imaging, including HTAN, HuBMAP, the BRAIN Initiative - Cell Census Network (BICCN), and the Kidney Precision Medicine Project (KPMP).

We asked those interested in attending to submit an application briefly describing: (1) an imaging-based computational challenge relevant to the cancer community, shared across multiple imaging modalities, with specific questions to address in a hackathon; (2) prior relevant work (their own or from the literature); and (3) data required to address the challenge, their availability, and whether they include any necessary annotated ground truth. The applications highlighted active areas of interest within the field. We grouped challenges described in the applications into four broad categories corresponding to stages in a canonical image processing workflow: (1) image registration and quality control; (2) segmentation of cells and subcellular structures; (3) downstream analyses, including cell type calling, cell tracking, and the discovery of spatial patterns; and (4) visualization and the integration of individual image processing steps into a larger automated pipeline. We organized the talks into sessions reflecting these canonical stages (Figure. 1a). Additionally, we invited three keynote speakers whose work intersected these stages and whose organizations were actively engaged in biomedical image acquisition and analysis: Drs. Susanne Rafelski (Allen Institute for Cell Science), Juan Caicedo (Broad Institute), and Matthew Cai (The Chan Zuckerberg Initiative).

**Figure 1:**
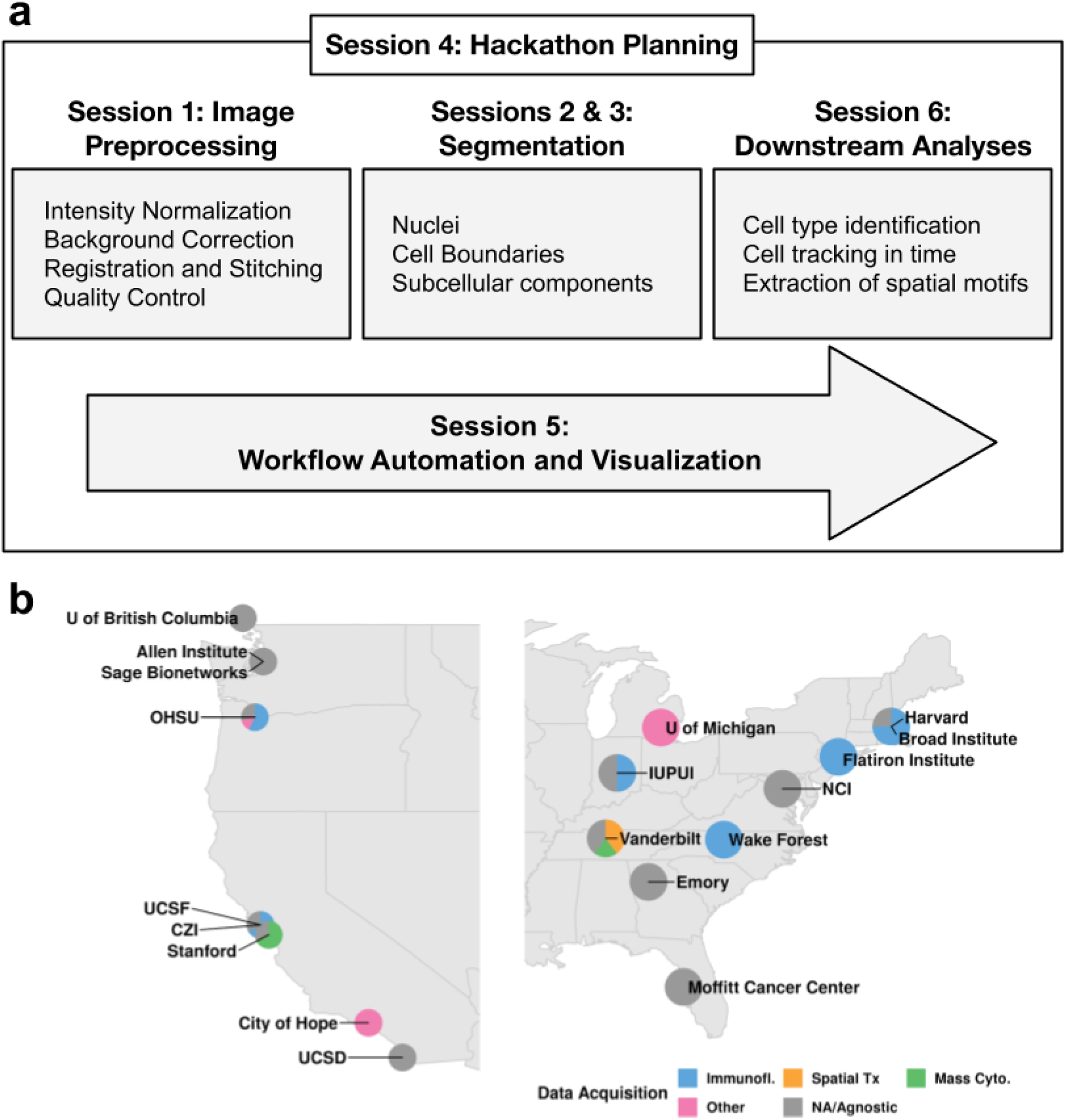
Image analysis workshop structure and participation. **a**. The workshop agenda followed the steps of a canonical image processing workflow. Each box highlights the most prominent topics covered during each session. **b**. Institutes and data acquisition technologies represented by the workshop participants. Technologies marked “other” encompass electron microscopy and radiology, while “NA/Agnostic” refers to computational labs that don’t generate data.

The explicit goal of the workshop was the generation of 2–3 page summaries describing each potential challenge. Prior to the workshop, attendees were provided with a template for these summaries, which included the same information solicited in the application (challenge idea, prior relevant work, and required data), as well as whether the challenge was best addressed collaboratively or competitively and, in either case, how to evaluate success. Example summaries representative of each of the four canonical stages were drafted based on several of the applications.

#### 2.2.2 Workshop implementation and management

The workshop took place Jan 19–20, 2020 at the Institute for Systems Biology (ISB) in Seattle, WA. The 46 attendees represented 12 institutions active in CSBC/PS-ON, as well as related consortia including HTAN, HuBMAP, the Imaging Data Commons (IDC), and KPMP (Table 1). Attendees were geographically distributed and came from laboratories employing a range of imaging modalities (Figure. 1b). They were predominantly early stage investigators (6 assistant professors and 1 associate professor), trainees (12 graduate students and 3 postdocs), and staff scientists (18). The size of the workshop fostered engagement and more than half of the attendees (25 of the 46) gave presentations. With the exception of the keynote speakers, we requested that all presenters structure their talks around one or more specific image analysis questions that could be expanded into a hackathon challenge. Each speaker was given a 20-minute time slot and encouraged to include the information previously described in the example summaries. To facilitate further discussion of these challenges after the talks, we scheduled ample additional time and allocated space for informal conversations.

**Table 1:** Workshop and hackathon attendee summary.

| Institute | Consortium | Workshop Attendees | Hackathon Attendees | Student | Postdoc | Staff Scientist | Faculty | Other |
| --- | --- | --- | --- | --- | --- | --- | --- | --- |
| Allen Institute | Unaffiliated | 3 | 2 | 0 | 0 | 2 | 0 | 2 |
| ASU | Unaffiliated | 0 | 1 | 1 | 0 | 0 | 0 | 0 |
| Broad Institute | Unaffiliated | 1 | 0 | 0 | 0 | 1 | 0 | 0 |
| City of Hope | CSBC | 1 | 1 | 0 | 0 | 2 | 0 | 0 |
| CZI | Unaffiliated | 1 | 0 | 0 | 0 | 1 | 0 | 0 |
| Emory | HTAN | 1 | 1 | 0 | 0 | 0 | 1 | 0 |
| Harvard | CSBC, HTAN | 2 | 5 | 1 | 1 | 3 | 0 | 0 |
| ISB | HTAN, IDC | 4 | 0 | 0 | 0 | 4 | 0 | 0 |
| IU/IUPUI | CSBC, KPMP, PSON | 2 | 5 | 4 | 1 | 0 | 1 | 0 |
| Mark III | Unaffiliated | 1 | 0 | 0 | 0 | 0 | 0 | 1 |
| Michigan | Unaffiliated | 1 | 0 | 0 | 0 | 0 | 0 | 1 |
| Moffitt | CSBC | 1 | 1 | 0 | 0 | 1 | 0 | 0 |
| NCI | CSBC, PS-ON | 2 | 0 | 0 | 0 | 0 | 0 | 2 |
| OHSU | CSBC, HTAN | 7 | 6 | 5 | 0 | 2 | 1 | 0 |
| QMUL | CSBC | 0 | 1 | 1 | 0 | 0 | 0 | 0 |
| Sage | CSBC, PS-ON | 1 | 1 | 0 | 0 | 1 | 0 | 0 |
| Simons Foundation | Unaffiliated | 1 | 1 | 0 | 1 | 0 | 0 | 0 |
| Stanford | CSBC, PS-ON | 5 | 2 | 0 | 2 | 1 | 0 | 0 |
| Susan G. Komen | Unaffiliated | 0 | 1 | 0 | 0 | 1 | 0 | 0 |
| UBC | Unaffiliated | 1 | 1 | 1 | 0 | 0 | 0 | 0 |
| UCSD | CSBC | 1 | 1 | 1 | 0 | 0 | 0 | 0 |
| UCSF | CSBC | 1 | 0 | 0 | 1 | 0 | 0 | 0 |
| Vanderbilt | CSBC, HTAN, HuBMAP | 8 | 19 | 9 | 2 | 6 | 3 | 1 |
| Wake Forest | CSBC | 1 | 0 | 0 | 0 | 0 | 1 | 0 |
| Weill Cornell | PSON | 0 | 1 | 1 | 0 | 0 | 0 | 0 |
| Total |  | 46 | 50 | 24 | 8 | 25 | 7 | 7 |

The workshop culminated in seven challenge ideas spanning image registration, automated marker gating, quantification of epithelial polarity in organoids, cell segmentation, quantification of PD1 asymmetry in immune cells, rare cell type identification in noisy data, and cell type inference from morphology and spatial distribution (Table 2). These ideas captured open research questions in the imaging field, though the questions continued to evolve through selection for and execution at the hackathon.

**Table 2:** Image analysis challenges nominated by workshop participants. Bolded challenges were selected for the hackathon.

| Challenge | Problem Statement | Goal |
| --- | --- | --- |
| **Image registration** | Deformations in tissue that arise in iterative staining approaches (e.g., CyCIF, 4i, IHC, MiFish) complicate alignment. | **Align images in a “z” stack (e.g., across rounds of staining or serial sections) to maximize the correlation of nuclei stains between consecutive images in the stack.** |
| Automated marker gating | Sample-sample variation in marker intensity owing to batch- and tissue-specific effects complicates efforts to compare markers in multiplex images across tissues and time points. | Automatically and consistently “gate” markers across tissues (i.e., to differentiate between expressed and unexpressed levels of marker expression). Predictions will be compared to manually-curated gating. |
| Quantification of epithelial polarity in organoids | Cells in organoids self organize and become polarized: specific proteins and organelles are oriented towards the center of the gland, whereas other cell components are expressed at the periphery. De-polarization may be associated with cancer. | Define a metric of organoid polarity and obtain values of this metric for each organoid in the provided data. |
| **Cell segmentation** | Developing a pipeline to segment nuclei and other cellular components (cell boundaries, subcellular structures, etc.) has traditionally been slow and incremental. | **Develop general guidelines for successfully optimizing segmentation algorithms via a competition. Teams will attempt to develop optimized segmentation algorithms for four different segmentation tasks within the hackathon.** |
| **Quantification of PD1 asymmetry in immune cells** | PD1 localization is asymmetric on a subset of cells during T-cell–tumor cell interactions and this asymmetry is thought to be related to T-cell activation. | **Develop a suitable metric and analytical pipeline for measuring the amount of polarization of membrane-bound markers (e.g. PD1).** |
| **Rare cell type identification in noisy data** | Known biology dictates the existence of certain rare cell subpopulations, but standard clustering methods fail to detect these robustly. | **Given a cell type definition (specified as marker expression), accurately find all cells of that type.** |
| **Cell type inference from morphology and spatial distribution** | Morphological characteristics may be relevant to discerning cell types but have not yet been used in cell type calling. | **Predict cell types (defined by characteristic marker expression) using morphological features (area, perimeter, eccentricity, etc.) and cell-cell proximity information (encoded as an undirected graph).** |

### 2.3 Hackathon

#### 2.3.1 Pre-hackathon planning

##### Selection of challenges

Given the large number of challenge ideas (Table 2) relative to the number of attendees we could support (∼40), we decided to limit the number of hackathon challenges. To ensure the attendees would be invested in the final set of selected challenges, we asked hackathon registrants to vote on their top three choices through a web-based form, which also captured additional information, such as their affiliated institutions, position held, and any special compute resource requests. This allowed for the assessment of both the interest in each of the proposed challenges and the willingness to contribute to different challenges in case a challenge had insufficient overall interest (less than three first-choice votes). Five of the seven challenges had sufficient interest and were selected for the hackathon (Table 2), with the Segmentation challenge garnering the largest proportion of interest (10 of 33 first-choice votes). Each challenge was assigned a champion, who ensured requisite data were transferred to Vanderbilt prior to the hackathon and who acted as a scientific advisor and hackathon liaison to the teams. Champions included the hackathon organizers, as well as Eliot McKinley (Vanderbilt University) and Seth Winfree (Indiana University) for the Segmentation Challenge.

##### Computational support

Through a partnership with Mark III Systems, Core Scientific provided access to an NVIDIA DGX2 containing 16 NVIDIA 32GB V100 GPUs and 4 × 24 core CPUs and 1 TB storage web-accessible from their datacenter in Dalton, GA; these resources were distributed among the participating groups. Mark III Systems also provided administrative support by preparing JupyterLab environments with software requested by attendees during registration, as well as on-site assistance to manage the environments. All data to be used for each of the challenges (∼500 GB total) was preloaded onto storage accessible by each of the compute instances. Notably, the collection and assembly of these large datasets from different sources and their deposition onto the different devices was a relatively time-consuming process, taking many hours of effort due to network latency and throughput limitations. As a backup, ten external 1 TB SSD hard drives were also preloaded with the full datasets, proving invaluable for the hackathon, due to the inability of several groups to use their preferred software tools within the prepared compute environments and allowing those groups to work locally without incurring the time-consuming overhead of downloading the data themselves.

Registrants were from over twenty different institutions (Table 1), with multiple from Vanderbilt and Vanderbilt University Medical Center (19), Oregon Health & Science University (6), Indiana University (including IUPUI and the School of Medicine, 5), Harvard University and Harvard Medical School (5), Stanford (2), and the Allen Institute for Cell Science (2). Trainees comprised a larger percentage of hackathon attendees (60%; 30 of 50) than of workshop attendees (24%; 11 of 46).

#### 2.3.2 Hackathon execution

The hackathon was hosted March 4–6, 2020 by Vanderbilt University to address the five selected challenges (Table 2). The Segmentation Challenge was set up in a competitive format and the (13) participating individuals were divided into three teams of three to five participants each. The remaining four challenges each had a team consisting of at least five participants. Work toward addressing the challenges was performed over two and a half days, coordinated by champions for each challenge and culminated with each team presenting their selected challenge and the solution they developed; this provided immediate feedback and guidance on future steps from the image analysis community.

## 3 Results

In this section, we provide an overview of analyses performed by the participants of every challenge. The code of all prototype solutions is publicly available on GitHub (Table 3), but additional work is required to generate well-documented easy-to-use software modules from the initial codebase. We conclude by summarizing the findings of each challenge in the context of the overall image analysis workflow and highlight notable gaps that we intend to address in future hackathons.

**Table 3:** Location of the code repositories for every challenge.

| Challenge | Code |
| --- | --- |
| Image Registration | https://github.com/IAWG-CSBC-PSON/registration-challenge,<br>https://github.com/jvizcar/MultiplexImagingRegistration |
| Segmentation | https://github.com/IAWG-CSBC-PSON/Segmentation-challenge |
| PD1 Asymmetry | https://github.com/IAWG-CSBC-PSON/pd1-asymmetry |
| Rare Cell Types | https://github.com/IAWG-CSBC-PSON/rare-cell |
| Cell Morpho-Typing | https://github.com/IAWG-CSBC-PSON/morpho-type,<br>https://github.com/jvizcar/SageMultiplexInteractors |

### 3.1 Image registration challenge

Iterative staining-based assays (e.g., CyCIF) involve successive rounds of staining with a small number of antibodies (∼4) conjugated to different fluorophores. These individual images need to be registered into a single composite, a task that often leverages a nuclear stain common across rounds (e.g., DAPI). For lower dimensional assays that can not rely on a shared stain across rounds, we explored registering images via auto-fluorescence.

During the hackathon, we observed that we could register a “moving” image in one round to a “target” image in a second round using the green channel (Alexa Fluor 488)—despite the fact that the fluorophore was conjugated to a different protein in each of the rounds (e.g., PCNA in round 1, CK5 in round 2, aSMA in round 3, etc.). We hypothesized that registration was exploiting the strong background fluorescence of the green dye. We provided further evidence for this hypothesis in additional analyses following the hackathon. In this follow-up work, we eliminated the possibility that registration was leveraging spatially overlapping signal between the different proteins in the two rounds by aligning the moving image to a target image consisting of only the background in the green channel. This often approached the resolution of the alignment obtainable using the DAPI channel shared across rounds in a healthy tonsil tissue, a healthy breast tissue, and a breast cancer cell line sample (Figure 2). Further, with one exception in the breast tissue, the background-derived registration aligned images with a resolution likely sufficient to associate cellular markers across rounds (i.e., that of a typical eukaryotic cell, ∼10**µ**m).

**Figure 2:**
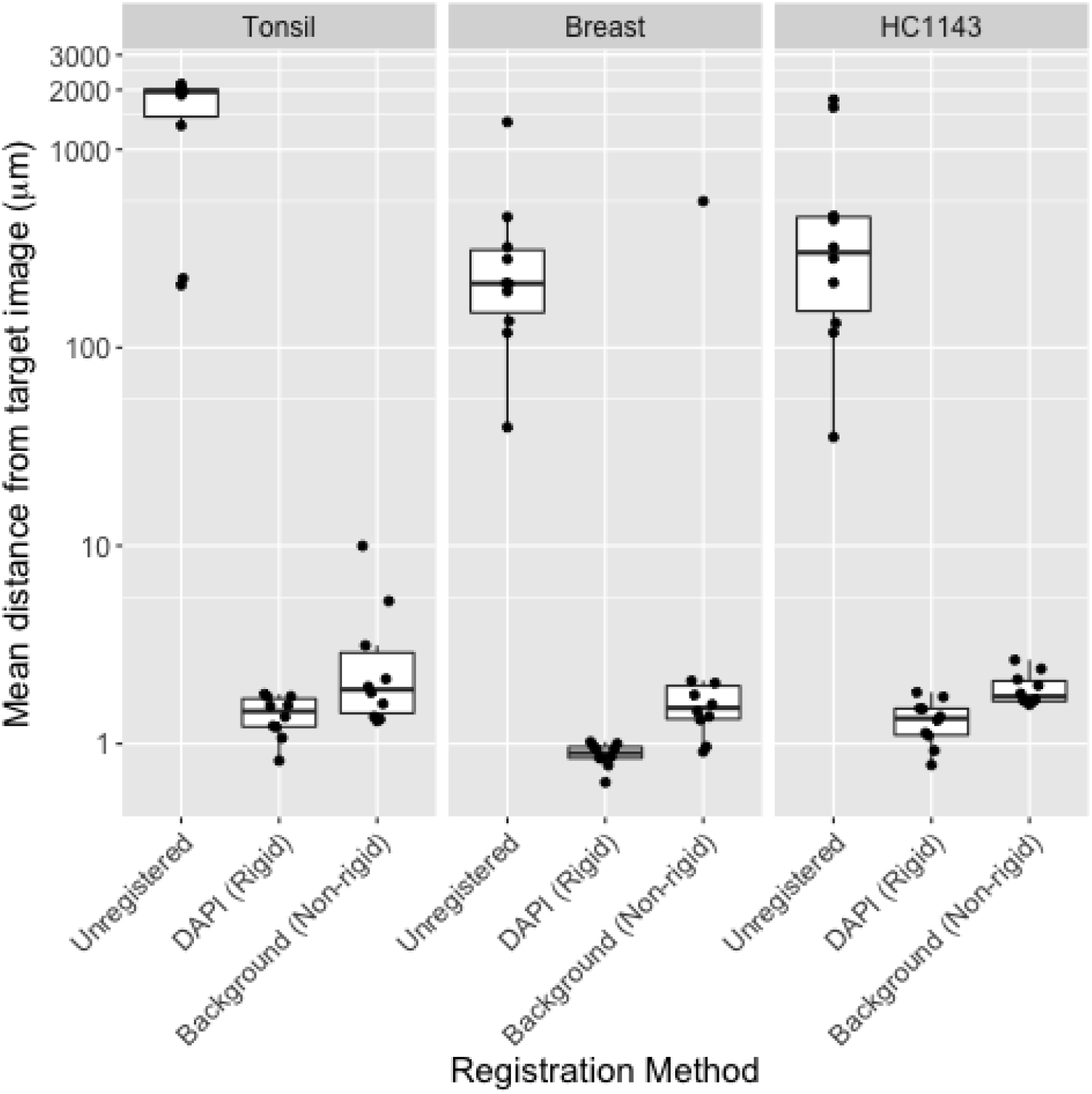
Image registration using background signal is similar to that obtained using DAPI. Mean distance from round 1 image (R1; y axis) to images in all other rounds (*n*=10) before registration (“Unregistered”), following rigid registration using the DAPI channel [“DAPI (Rigid)”], or following a non-rigid registration using the background channel [“Background (Non-rigid)”] in tissue microarray images from healthy tonsil tissue (left), healthy breast tissue (center), or HC1143 breast cancer cell line (right). Distance is calculated by defining AKAZE (Alcantarilla et al., 2013) keypoints in both images, matching the keypoints across images, summarizing the distance between matched keypoints using target registration error (TRE) (Maurer et al., 1997), and scaling TRE (whose denominator is the diagonal of the image in pixels) by the length of the image diagonal in microns. Non-rigid registration was performed using SimpleElastix (Marstal et al., 2016); rigid registration was performed using AKAZE-based keypoint matching in OpenCV (Bradski, 2000).

### 3.2 Segmentation challenge

Developing a pipeline to segment nuclei and other cellular components (cell boundaries, subcellular structures, etc.) has traditionally been slow and incremental. Several tools are available that can apply the multiple steps required for optimum segmentation, including Cell Profiler, ImageJ, Matlab, and the Cell and Structure Segmenter developed at the Allen Institute for Cell Science. The time required to develop these pipelines is a hindrance to extracting biologically useful information, yet there are very few guidelines to facilitate the process. Thus, a main goal of this challenge was to identify general guidelines for successfully optimizing segmentation algorithms. The challenge was structured as a competition, where teams competed to perform various segmentation tasks on several large datasets with a focus on balancing segmentation accuracy with the ability to process all the data. Only laptops brought to the hackathon or the compute environments made available to all attendees were allowed, as the use of an external high-performance computing environment would have provided an unfair advantage. The Allen Cell and Structure Segmenter (Chen et al., 2020) was pre-installed within the compute environments provided to all teams.

Five different cancer-relevant segmentation tasks were identified, representing a range of different features, with all requiring initial nuclear segmentation (Table 4). Each task had an associated dataset consisting of fluorescence microscopy images (color images of typical histopathological evaluation using H&E counterstaining were not considered). Datasets were provided by members of the IAWG or were selected from publicly available resources. Each dataset contained many visual fields of information, some stitched into image montages and some containing multiple channels obtained from the same field of view. The total size of all image data combined was over 250 GB, making the volume of data for processing a significant hurdle.

**Table 4:**
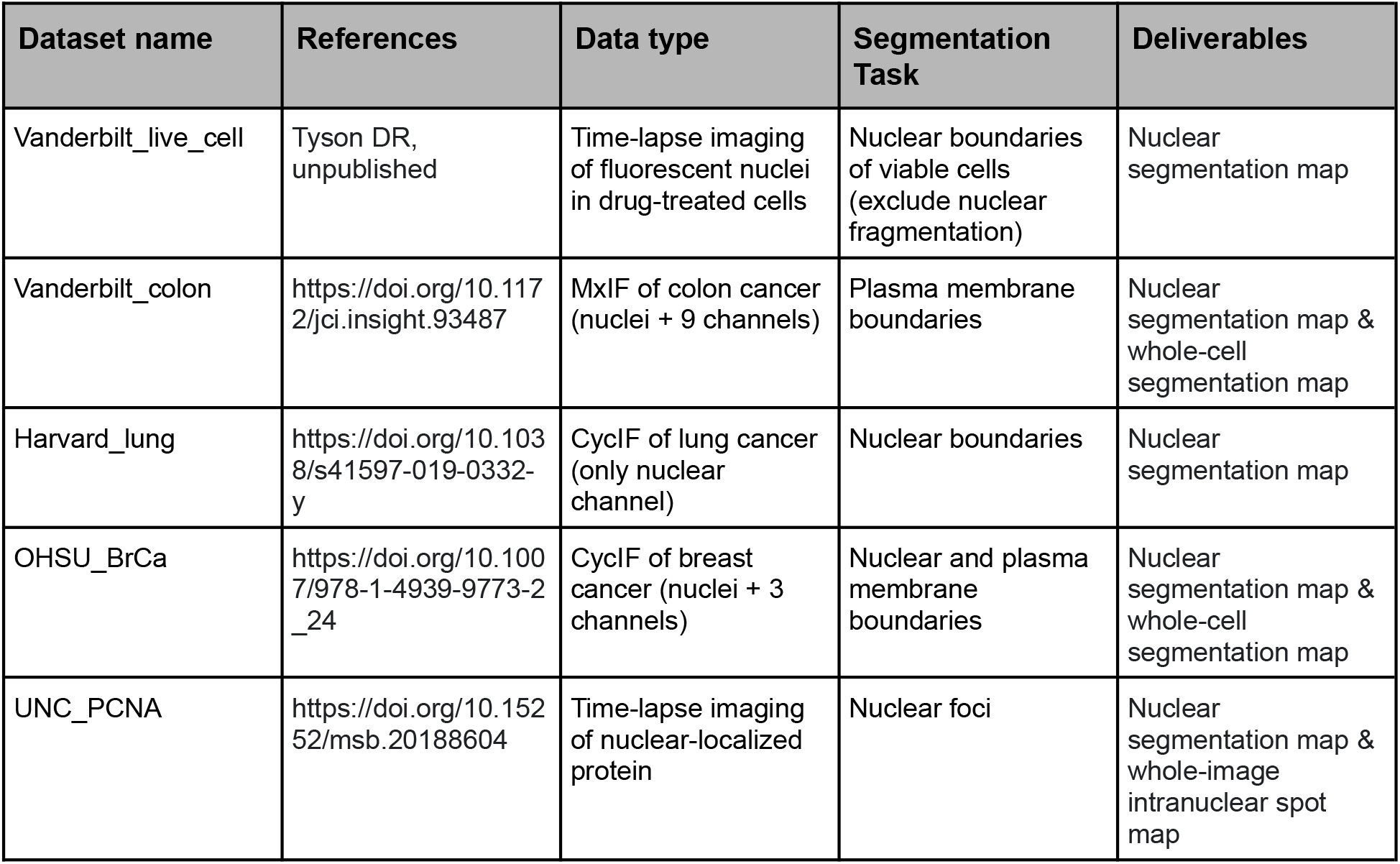
An overview of datasets contributed to the segmentation challenge.

At the onset of the hackathon, three teams were formed by self-association among the attendees, resulting in two teams of five members and one team of three members. To assess the quality of each team’s segmentation results a small Java program was written to calculate F1-scores on segmentation results as compared to previously generated ground-truth labeled images (SegmentationAnalyzer, code available at https://github.com/IAWG-CSBC-PSON).

#### 3.2.1 Segmentation Team 1

Team 1 used a KNIME-based workflow and ImageJ processing (Dietz et al., 2020) for segmentation of the MxIF and CycIF data; however, this pipeline could not be deployed on the provided compute server and was instead run on the team’s own computers. The live-cell data was segmented using the Scientific Python (SciPy) Multidimensional image processing (ndimage) package for segmentation. Overall Team 1 processed examples from nearly all datasets (4 out of 5) of the total dataset and on the directly comparable dataset (Vanderbilt live-cell images), obtaining a pixel-wise F1-score of 0.33.

#### 3.2.2. Segmentation Team 2

Team 2 leveraged several pre-existing tools, including CellDissect (Kesler et al., 2019) and DeepCell (Valen et al., 2016) for nuclear segmentation and made use of algorithms for intensity gradient detection and K-means clustering to detect the colonic epithelial cell boundaries. Team 2 processed examples from all the datasets and for the directly comparable dataset achieved a pixel-wise F1-score of 0.32, which, in combination with completing examples from all datasets (5 out of 5), gave them an advantage over Team 1.

#### 3.2.3 Segmentation Team 3

Team 3 used Python-based processing, including the Allen Cell and Structure Segmenter code (Chen et al., 2020) and focused on the Vanderbilt live-cell image dataset. They chose not to submit their results for the competition, preferring to use the time simply as a learning experience.

#### 3.2.4 Segmentation Challenge Outcome Summary

The volume of data and the multiple data types to be segmented posed significant barriers to rapidly developing high-performing algorithms. Based on the relatively low F1 scores (<0.35), it is clear that, even with contributions from experienced individuals who have published image segmentation pipelines, finding optimum solutions to specific image segmentation objectives remains a significant challenge, requiring much more time than was available during the hackathon to develop efficient solutions.

### 3.3 Quantitation of immune checkpoint markers (PD1) asymmetry in activated immune cells

PD-1 and PD-L1 represent perhaps the most well-known receptor-ligand pair that is targeted by immune checkpoint therapies, with a number of FDA approved drugs (mostly monoclonal antibodies) designed against both proteins. Transmembrane PD-L1 is most commonly expressed on tissue and stromal cells, while PD-1 can be seen in the T and B cells exposed to the antigen. If a PD-1 positive T cell is triggered by a cell expressing PD-L1, the PD-1 expression tends to spatially co-cluster with the activated T-cell receptor (TCR) and bring the phosphatase to the intracellular part of the TCR complex. This inhibits signal transduction from the activated TCR complex and corresponding downstream events associated with normal T cell response. Microscopically, these molecular processes are manifested in co-polarization (co-clustering) of PD-1 and TCR distribution in the immune cells engaged in interaction with their microenvironment. Being able to quantify the incidence of asymmetric distribution of PD-1 and TCR and of related molecules will therefore enable deeper understanding of factors involved in the normal immune response to cancer and provide insight into why this response may fail in specific scenarios.

Participants were asked to quantitate PD-1 clustering in highly–multiplexed CODEX images (Goltsev et al., 2018) acquired from lymph nodes of mice challenged with metastatic melanoma cell lines. The multiplexed images (∼72 channels, 7 z-planes, ∼25 frames, ∼3Gb per image) were segmented, with the center and the outline of the best focal plane for each cell saved in the text format, alongside measurements for roughly 50 different markers. Using the segmentation output, participants constructed image patches centered around individual cells and trained a Variational Autoencoder (VAE) (Kingma and Welling, 2014) to study the underlying latent space representation of the data. The spatial distribution of PD-1 in a population of cells is expected to follow a smooth and continuous distribution. Consistent with that, we found that the UMAP embedding of the latent VAE space is also continuous without any obvious cluster structure (Figure 3a). Mapping the mean PD-1 signal per cell onto the embedding, we observed two groups of cells with significantly elevated PD-1 intensities, indicating that the autoencoder is picking up on and encoding not just the spatial distribution, but also signal intensity in its latent space. We further subdivided the latent space into evenly spaced regions and plotted the average PD-1 distribution per region, revealing that the regions indeed display distinct spatial arrangements of PD-1 (Figure 3b). For example, regions 12 and 14 contain cells with a roughly even distribution of PD-1 across the entire cell membrane. In contrast, regions 4 and 5 correspond to highly polarized PD-1 distributions at the top-left or bottom-right of the cells, respectively. In the future, this method could be simplified by taking advantage of the rotational symmetry in the data, e.g., by rotating all cells such that the brightest pixel is always placed at the center-top part of the image. Alternatively, one can take advantage of the recently proposed Multi-Encoder VAE (Ternes et al., 2021) to extract transform-invariant features.

**Figure 3:**
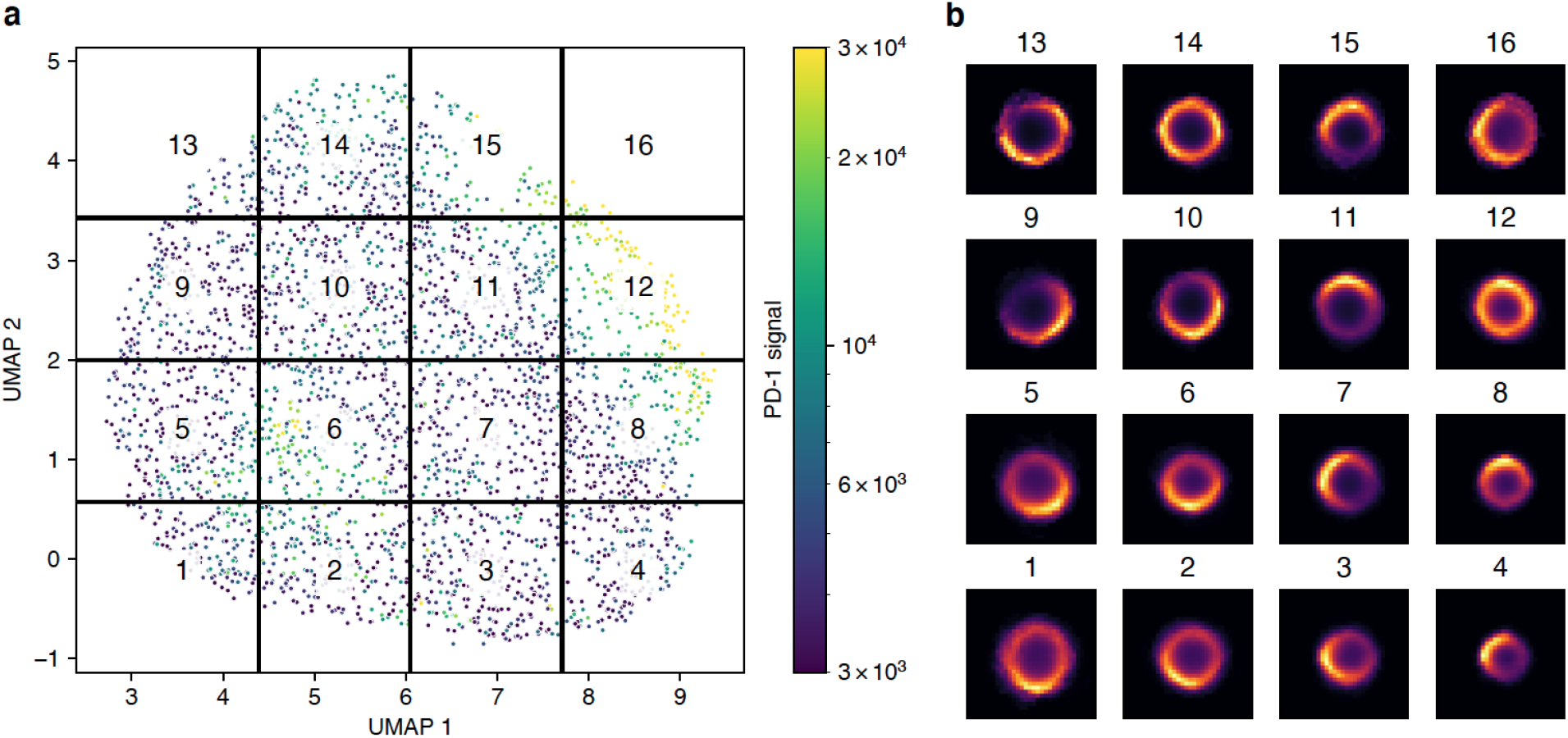
Spatial distribution of PD-1 classified using a variational autoencoder. **a)** UMAP embedding of the latent space produced by the autoencoder. Cropped images of cells stained for PD-1 were used to train a convolutional autoencoder with a bottleneck size of eight. The latent representation of all cells was further reduced using UMAP. Each point in the plot represents a single cell colored by the mean intensity of the PD-1 signal. The UMAP space was evenly divided into 16 regions. **b)** Mean spatial signal of PD-1 across all cells contained within each region.

### 3.4 Rare cell type identification in “noisy” multiplexed in situ cytometry data

Differentiation cascades in mouse immune organs are sufficiently well dissected by flow cytometric methods, yet the spatial architecture of immune cell development is not well understood. At the same time, the “curse of dimensionality” as well as noise in single-cell segmentation and quantitation preclude efficient cell type identification by clustering of *in situ* cytometric data. We often know that a particular rare cell type exists in an organ, and may be visible in the image, yet it is hard to unambiguously identify these cells by empirically-chosen machine learning techniques. It is therefore important to be able to select the best normalization, pre-processing, and clustering methods that result in proper identification of rare cell types when applied to the given data.

Participants attempted to identify specific rare cell types in multiplexed (∼50 markers) mouse thymus imaging data. These data were presented to participants in the derived form of a cell-by-feature dataframe, where each cell was represented by a row vector of mean marker instensities. To normalize the raw data, vertical (feature-based) and horizontal (cell-based) strategies were considered, both independently and in combination. In addition to boilerplate standardization, the participants also attempted to leverage known mutually-exclusive expression patterns of some marker pairs to derive normalization factors for markers where the mutually-exclusive expression assumption was met (Chang et al., 2020). Using data normalized by each approach, the participants then used a battery of automated single-cell phenotyping approaches in an attempt to identify the rare immune cells populations. The results of cell type identification were scored against manually-gated ground truth labels to characterize the performance trade-offs of taking each combination of pre-processing and analysis approach. In the mouse thymus dataset, composed primarily of hierarchically-structured immune cell phenotypes, the participants found that the approach combining first vertical then horizontal standardization was necessary to identify the rare cell types by any subsequent phenotyping method, e.g. *k*-means clustering and X-shift (Samusik et al., 2016).

### 3.5 Inferring Cell Type from Morphological Features and Spatial Distribution Data

By visual inspection, one can readily appreciate the difference in shape and size between various cell types, as well as mesoscale structures defined by the arrangement of cells within tissues (Figure 4a). Using publicly available CyCIF data from three lung cancer specimens (Rashid et al., 2019), we formulated a challenge focused on inferring the type of a cell directly from its morphological features and the spatial distribution of its neighbors. The participants were asked to design and train a cell type predictor that accepts as input 1) a vector of morphological features such as area, perimeter, eccentricity, etc., and 2) cell-cell proximity information encoded as an undirected graph. The prediction task was formulated as a three-class classification problem with ground truth labels derived from marker expression, based on the following known marker to cell type associations: CD45 - immune cells; Keratin - tumor cells; and alpha-SMA - stroma. The morphological features were computed using the Python-based package scikit-image (van der Walt et al., 2014), and each cell was annotated with its five closest neighbors, based on Euclidean distance in the image coordinate space.

**Figure 4:**
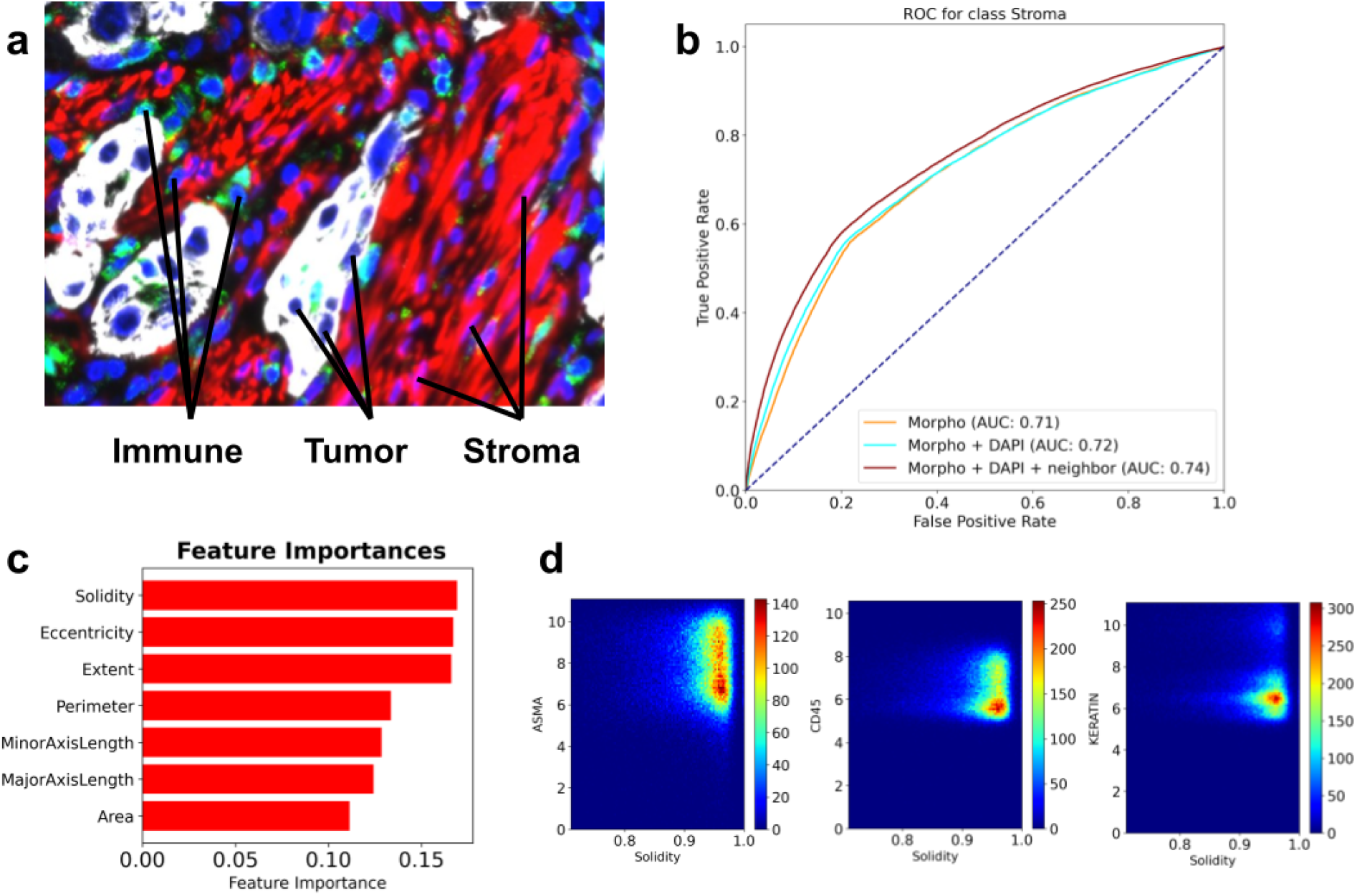
A summary of the *morphotype* challenge. **a)** A representative image of a lung cancer specimen, stained for Keratin (white), CD45 (green), IBA1 (cyan) and alpha-SMA (red). **b)** ROC curves associated with predictors trained to recognize stromal cells from morphological features only (yellow), morphological features and the intensity of the DAPI stain (cyan), and all of the above computed for the index cell and its closest five neighbors (brown). **c)** Top features identified through feature importance scores computed by gradient boosted random forests. **d)** Density scatter plots showing how solidity varies with markers of stromal (alpha-SMA), immune (CD45) and tumor (Keratin) cells.

Challenge participants decomposed the problem into a collection of three “one cell type vs. the other two” classification tasks and trained gradient-boosted random forest models (Friedman, 2001) for each binary task. Metaparameter tuning was performed using grid search and five-fold cross-validation over the training set, and the final models were evaluated via leave-one-image-out cross-validation. The participants found that morphological features carried modest signal predictive of cell type, with area under the ROC curve (AUC) being in the 0.7-0.8 range for all models (Figure 4b). Very minor improvements in performance were achieved by incorporating morphological features of direct neighbors and intensity of the DAPI channel (a proxy for the size of the nuclei). Further inspection of the feature importance scores revealed that solidity (the ratio of an area to its convex hull), eccentricity (the ratio of the focal distance to the major axis length) and extent (ratio of pixels in the region to pixels in the total bounding box) were most informative for distinguishing between tumor, immune and stromal cells (Figure 4c). Direct inspection of correlation between solidity and the three cell type markers (CD45, Keratin and alpha-SMA) revealed that high values of solidity were observed in stromal cells and, to a lesser extent, immune cells (Figure 4d). Based on the challenge outcome, the participants concluded that a cell type caller based solely on morphological features is unlikely to be sufficiently accurate, and future improvements are expected from combining morphological information with marker expression patterns.

### 3.6 Image processing workflow

Our hackathon challenges have a direct correspondence to individual steps in a canonical image processing pipeline. Because these steps are executed sequentially, an important property of the workflow is that upstream processing, such as image registration, has a direct impact on the quality of inputs for segmentation, which in turn affects the quality of inputs for downstream analyses like cell type calling. It is therefore imperative to establish best practices to ensure that upstream errors do not get amplified in downstream processing. Our final result is an outline of what we envision these best practices to be, based on lessons learned during the hackathon challenges.

The original goal of the registration challenge was to mitigate physical tissue deformations (e.g., tearing or folding) through use of non-rigid registration approaches that “correct” them. However, in applying them to align images based on background fluorescence, we observed that these non-rigid approaches can greatly distort the moving image (e.g., shrinking it dramatically) when it differs greatly from the target image. As such, these methods should be applied with considerable care, possibly constrained or assisted by deep learning (Fu et al., 2020; Haskins et al., 2020).

The segmentation challenge highlighted that the difficulty of a segmentation task is driven primarily by the imaging modality and the tissue being imaged: cells in culture were substantially easier to segment than tissue images, particularly those with densely-packed cells (e.g., Vanderbilt_colon; Table 4). We hypothesize that training tissue-specific models will play a bigger role in accurate segmentation than the underlying machine learning methodology. Emerging studies also show that data augmentation and the inclusion of a nuclear envelope stain, such as Lamin, can substantially improve segmentation accuracy in a method-agnostic way (Yapp et al., 2021).

Downstream cell type calling still heavily relies on prior knowledge about the association of cell types with certain markers. The prior knowledge can help mitigate errors from upstream processing by ensuring that marker expression aligns with known biology, but extra care must be taken to ensure that segmentation artifacts are not misinterpreted as novel or rare cell subpopulations. Our future hackathon efforts will focus on evaluating the interplay between segmentation and cell type calling, as well as systematically assessing how degradation in upstream performance impacts the quality of downstream analyses. We also expect that future cell type calling will work directly at the pixel level, allowing methods to take full advantage of the spatial distribution in signal intensities (as demonstrated by our PD-1 polarity challenge) and the power of existing deep learning architectures (He et al., 2021). We will formalize this expectation into a pixel-level cell type prediction challenge in a subsequent hackathon.

## 4 Discussion

While most hackathons involve pre-hackathon planning activities, our hosting of a two-day workshop to formalize the challenges was a novel aspect that, to the best of our knowledge, has not been attempted before. The workshop allowed democratization of the challenge questions by soliciting input from the image community as a whole and narrowing those down based on the participant interest. As a result, the participants were able to address a diverse range of specific questions that required substantially different approaches, while exchanging ideas on issues common to all challenges, such as the need to wrangle large, complex image datasets, train machine learning models, and visualize the results. This was a particularly timely experience, as increasingly-sophisticated imaging methods become more widespread in cancer research, and many groups grapple with the best way to analyze data that is rapidly growing in both size and complexity.

Our original intent was to have more established investigators frame challenges in the field during the workshop. However, attendees at both the workshop and hackathon were mostly early-stage investigators and trainees and most attended both events. This had important advantages: hackathon attendees were invested in the challenges that they themselves had previously defined during the workshop and they had already developed a rapport with their team members. Having two face-to-face meetings with a small and highly overlapping group of participants also improved the likelihood of forming post-event collaborations. Indeed, the PD1 asymmetry team continued their efforts after the hackathon. Hence, we feel the preliminary workshop was critical in realizing the hackathon outcomes (including scientific results and collaborations) and that replicating either experience virtually would have been difficult. Consistently, all eight post-hackathon survey respondents who also attended the workshop reported that their participation in the first event was useful in preparing for the second.

To assess the impact of the workshop and the hackathon, we conducted a survey with questions about what the participants found most valuable and what could be improved. Feedback from the participants confirmed many positive aspects of hands-on working meetings described in the literature (Groen and Calderhead, 2015; Huppenkothen et al., 2018). Specifically, the meetings provided a good networking opportunity for scientists from different labs and with different areas of expertise, allowing researchers to establish new collaborations and brainstorm ideas for future projects. As described previously, such opportunities are particularly important for early-career scientists, including postdocs and junior investigators (Groen and Calderhead, 2015).

The hackathon also provided a hands-on educational experience by bridging the gap between traditional courses, which take months to develop, and the rapidly shifting landscape of image analysis tools. By exposing the participants to real datasets with all their complexities, the hackathon was an immersive experience that fostered collaborative software development and an exchange of ideas that can be taken back to each lab’s day-to-day activities. All of the survey respondents expressed interest in participating in future image analysis hackathons.

Our community-based approach had several limitations. Previous hackathons highlighted the importance of having teams composed of participants with diverse backgrounds (Ferreira et al., 2019; Groen and Calderhead, 2015). Unfortunately, the vast majority of our participants were computational scientists; having more representation of experimental scientists among the teams would have likely increased the biological and clinical insight produced by individual challenges. Several groups had a range of experience levels that provided learning opportunities for more junior members, as well as the requisite background to ensure productivity in the short time window of the hackathon. The registration team reported benefitting from several expert practitioners, suggesting that future imaging-based hackathons should strive to ensure experts are embedded in each team. Counterintuitively, background diversity among the computational scientists was also a limitation; Python is widely considered to be the primary language for image analysis tasks, and participants with background in other programming languages (e.g., Java or R) felt at a disadvantage. We conjecture that the use of container technologies, such as Docker (Merkel, 2014) and Singularity (Kurtzer et al., 2017) in future hackathons can help with code execution across various compute environments and programming language preferences.

The hackathon exposed three primary technical hurdles likely to be pervasive in large-scale image analysis projects: (1) access to GPUs for more efficient computation; (2) scalable access to large datasets for generalizing trained models; and (3) access to ground truth for objective evaluation of those models. The likely performance advantage of GPUs would have been especially beneficial to the segmentation challenge, in which one team had inadequate time to analyze all datasets. Across all of the challenges, lack of “ground truth” labels made it difficult to effectively evaluate the solutions produced by the hackathon participants—an oft-occurring difficulty in biomedical image analysis, as producing a reference standard traditionally requires laborious, manual curation by pathologists (Willemink et al., 2020). To mitigate this, data contributors sometimes had to rely on orthogonal measurements to generate label approximations (e.g., using expression of protein markers to define cell types for the cell morphology challenge). Nevertheless, for the prototyping purposes of our hackathon, most teams found that CPUs were sufficient to assess their approaches against a few exemplar images and that those few images could be directly accessed from the portable hard drives we provided. This may serve as a model for future image analysis hackathons: limit the scope of proposed challenges to be practically addressable using local hardware with a scale of data that can be transferred across a network or distributed via external hard drives. Given the limited scale of such data, it would also be feasible to provide attendees early access to them *prior* to the hackathon to facilitate data exploration without time constraints of the event—an opportunity our participants regretted not having.

However, rather than limiting the scale of a future hackathon, we propose a more ambitious goal: address the above issues by conducting the hackathon in a cloud environment. A community-wide, shared, repository co-localized with compute infrastructure in the cloud would also facilitate the collaborative efforts that our hackathon showed to be both educationally and scientifically productive. Efficiently integrating image analysis with cloud resources remains a challenge, owing to the latency of data transfer and of remote interactive viewing. Integrated solutions for image storage, viewing, and analysis have been implemented on high-performance computing (HPC) clusters for pathological (H&E) images (Schüffler et al., 2021). Such approaches would need to be further extended to account for the order of magnitude greater storage requirements of multiplexed data and migration from an HPC environment to the cloud to further ease cross-institutional collaborations. Members of our community have already taken strides towards addressing these issues by developing Minerva Story (Hoffer et al., 2020) and the Cancer Digital Slide Archive (CDSA) (Gutman et al., 2013) for scalable visualization of highly multiplexed and H&E image data, respectively, and through MCMICRO, a workflow-based, configurable image analysis pipeline (Schapiro et al., 2021) that facilitates cloud-based computation by adopting containerization technologies, such as Docker and Singularity, and seamless access to heterogeneous resources, such as GPUs. Finally, recent advances in cell segmentation have demonstrated how ground truth datasets can be efficiently generated through cloud-based, crowd-sourced annotations that are verified, rather than generated *in toto*, by expert annotators (Greenwald et al., 2021). The difficult work remains in bridging scalable visualization platforms with analytical frameworks and assessing them on ground truth datasets scalably in the cloud. Nevertheless, even partially demonstrating this feasibility in a subsequent hackathon would advance future cross-institutional projects.

## Acknowledgments

We gratefully acknowledge funding support from the NCI grants U24CA209923 and U54 CA217450-02S1, which allowed for reimbursement of accommodation and travel expenses for approximately 40 participants to each event. We would like to thank Daniel Gallahan and Shannon Hughes for their insightful comments and suggestions, as well as Mark III Systems and Core Scientific for providing access to their cloud environment and for facilitating on-site administrative support.

